# Open-Access STRS Database Of Populations From The 1000 Genomes Project Using High Coverage Phase 3 Data

**DOI:** 10.1101/2021.09.06.459168

**Authors:** Tamara Soledad Frontanilla, Guilherme Valle-Silva, Jesus Ayala, Celso Teixeira Mendes-Junior

## Abstract

Accurate STR genotyping from next-generation sequencing (NGS) data has been challenging. Haplotype inference and phasing for STRs (HipSTR) was specifically developed to deal with genotyping errors and obtain reliable STR genotypes from whole-genome sequencing datasets. The objective of this investigation was to perform a comprehensive genotyping analysis of a set of STRs of broad forensic interest from the 1000 Genomes populations and release a reliable open-access STR database to the forensic genetics community. A set of 22 STR markers were analyzed using the CRAM files of the 1000 Genomes Project Phase 3 high-coverage (30x) dataset generated by the New York Genome Center (NYGC). HipSTR was used to call genotypes from 2,504 samples from 26 populations organized into five groups: African, East Asian, European, South Asian, and admixed American. The D21S11 marker could not be detected in the present study. Moreover, the Hardy-Weinberg equilibrium analysis, coupled with a comprehensive analysis of allele frequencies, revealed that HipSTR could not identify longer Penta E (and Penta D at a lesser extent) alleles. This issue is probably due to the limited length of sequencing reads available for genotype calling, resulting in heterozygote deficiency. Notwithstanding that, AMOVA, a clustering analysis using STRUCTURE, and a Principal Coordinates Analysis revealed a clear-cut separation between the four major ancestries sampled by the 1000 Genomes Consortium (AFR, EUR, EAS, SAS). Meanwhile, the AMOVA results corroborated previous reports that most of the variance is (97.12%) observed within populations. This set of analyses revealed that except for larger Penta D and Penta E alleles, allele frequencies and genotypes defined by HipSTR from the 1000 Genomes Project phase 3 data and offered as an open-access database are consistent and highly reliable.

## INTRODUCTION

Next-generation sequencing (NGS), massively parallel or deep sequencing, is a technology that enables the sequencing of millions of DNA fragments in parallel. NGS can deal with several regions or targets simultaneously, enabling the detection of variation sites or mutations in the genome. This technology has allowed the in-depth study of worldwide human genetic diversity for various purposes, including forensic human identification (1–3).

The 1000 Genomes Consortium is a worldwide collaboration that has produced an extensive catalog of human genetic variation. They sequenced whole genomes from 2,504 individuals from multiple populations derived from five groups: African, East Asian, European, South Asian, and admixed Americans (4). These data are freely available at the International Genome Sample Resource (IGSR) website to generate a variant call format file using a set of specific command lines (5). In 2015, they analyzed the genomes of all individuals using a combination of low-coverage whole-genome sequencing (WGS), deep exome sequencing, and dense microarray genotyping in project’s phase 3. They have described worldwide patterns of genomic diversity based on Single Nucleotide Polymorphisms (SNPs), indels, and structural variants (SVs), including deletions, insertions, duplications, inversions, and CNVs (copy-number variants). However, short tandem repeat (STR) markers were not analyzed and studied in depth (6).

STR markers are crucial in human identification due to the high polymorphism levels, being particularly useful for interpreting mixtures of biological samples. In addition to the issue of small-sized amplicons, genotyping these markers using NGS data is particularly difficult due to frequent alignment and stutter errors (7). The accurate genotyping from NGS data has been challenging because of their high sequencing error rates (8). A previous study managed to obtain and analyze STR data from the 1000 Genomes dataset using lobSTR (9). Since high coverage is mandatory for reliable STR genotype calling, a primary concern regarding that study is that the 1000 Genomes data available for lobSTR was generated using shallow sequencing coverage (2x – 6x), so it was severely susceptible to errors.

The New York Genome Center recently resequenced the 2,504 samples from 1000 Genomes Project Phase 3 panel with high (30x) coverage. They have also aligned the sequence data to GRCh38, and these publicly available alignments could be used to call STRs reliably (5, 10).

Some tools have already been developed to analyze STRs markers from NGS data, such as lobSTR (9), STRait Razor (11), toaSTR (12), HipSTR (13), and others. Each software uses different algorithms and flanking regions to capture STR reads.

Haplotype inference and phasing for STRs (HipSTR) was developed for calling microsatellites from Illumina FASTq files. It was designed to deal with genotyping errors and to obtain more robust STR genotypes. HipSTR accomplished this by learning locus-specific PCR stutter models using an EM algorithm, employing a specialized hidden Markov model to align reads to candidate alleles while accounting for STR artifacts, and using phased SNP haplotypes to genotype and phase STRs. These factors turn HipSTR into one of the most reliable tools for genotyping STRs from Illumina sequencing data (13).

Different from other tools, it can process hundreds of samples at once. Also, it enables the user to determine the set of STR markers to be analyzed, and the flanking regions used to capture them. In fact, HipSTR showed accurate genotype calling in previous studies. HipSTR accuracy was tested by comparing whole-genome sequencing calls from 118 samples to capillary electrophoresis data, resulting in a 98.8% consistency (13, 14). Recently, we performed a study comparing HipSTR with Strait Razor and toaSTR, and all three methods presented high allele calling accuracy (greater than 97%) (Valle-Silva, Frontanilla, et al., 2021, in preparation). Although data processing with HipSTR is more complex and requires bioinformatics knowledge and some nomenclature adjustments (detailed in Valle-Silva, Frontanilla et al. 2021, in preparation), HipSTR has shown to be the fastest and most appropriate tool to deal with larger datasets, including whole genomes.

This investigation aimed to perform a comprehensive genotyping analysis of a set of STRs of broad forensic interest from the 1000 Genomes populations and release a reliable open-access STR database to contribute to future forensic genetics studies.

## METHODOLOGY

### Genotype Calling

The CRAM files of 1000 Genomes Project Phase 3 resequenced high-coverage (30x) WGS data generated by the New York Genome Center (NYGC), available at https://www.internationalgenome.org/data-portal/data-collection/30x-grch38, were used. The sample size consisted of 2,504 individuals from 26 worldwide populations that compose five population groups: African (AFR), East Asian (EAS), European (EUR), South Asian (SAS), and admixed American (AMR)(4). All samples were sequenced with a paired-end approach (2 × 150pb), using the NovaSeq 6000 Sequencing System (Illumina, Inc.). The high-coverage data provides a more reliable opportunity to genotype STR markers. Thus, we have selected a set of 22 autosomal microsatellites commonly used in forensic practice: CSF1PO, D1S1656, D2S441, D2S1338, D3S1358, D5S818, D7S820, D8S1179, D10S1248, D12S391, D13S317, D16S539, D18S51, D19S433, D21S11, D22S1045, FGA, Penta D, Penta E, TH01, TPOX, and vWA. These 22 *loci* are autosomal markers that compose the PowerPlex^®^ Fusion System (Promega, Madison, WI).

We ran the HipSTR algorithm for each individual to genotype the 22 STRs based on the human reference genome GRCh38, using a BED file with the flanking regions available in the HipSTR repository (13). We applied the calling filter (15% stutter model) and a minimum of 8 reads to obtain more reliable genotypes. According to a binomial distribution, this minimum number of reads ensures (*p* > 0.99) that a homozygous genotype is called because of lack of variability at a given locus and not because the second allele was not sampled (Valle-Silva, Frontanilla, et al., 2021, submitted).

To perform genotype calling, we used the VCF output file produced by HipSTR taking three parameters into account: the reference allele of each marker, the period (i.e., the length of each STR motif), and the base pair differences (GB) in comparison with the reference allele. Nomenclature adjustments were made for D19S433, Penta D, Penta E, and vWA following (Valle-Silva, Frontanilla, et al. 2021, submitted) recommendations: removal of two repeat units from all D19S433 and vWA alleles, the inclusion of one repeat unit into all Penta D alleles, and removal of two nucleotides from all Penta E alleles. The total number of reads used to genotype each marker in each sample was also recorded.

### Statistical Analysis

Allele frequencies, Hardy-Weinberg equilibrium, and forensic parameters [Match Probability (MP), Power of Discrimination (PD), Power of Exclusion (PE), and Polymorphism Information Content (PIC)] were calculated for each population sample or each population group using the GenAlEx (15) and STRAF (16) software.

A Principal Coordinates Analysis (PCoA) using GenAlEx (15), Analysis of Molecular Variance (AMOVA) using Arlequin (17), and clustering analyses using STRUCTURE 2.3.4 (18) were used to explore the distribution of genetic diversity across populations of different ethnic backgrounds. The STRUCTURE analysis was performed for *k* ranging from 3 to 6, applying the correlated allele frequencies model, 100,000 burn-in steps followed by 100,000 Markov Chain Monte Carlo interactions, in 100 independent runs. The results from the runs with the largest “Estimated Ln Probability of Data” [LnP(D)] were selected and depicted in bar plots created with Distruct 1.1 (19).

We also compared the allele frequencies estimated from the 1000 Genomes dataset with STR data retrieved from the same five major population groups (African, European, East Asian, South Asian, and admixed American) that compose the SPSmart STR browser (Pop.STR) (20). For this purpose, allele frequencies of each STR for a given population group were compared between the two datasets using *F*_*ST*_ and an exact test of population differentiation based on genotype frequencies, using the Arlequin software (17). This comparison was made to verify the reliability of genotype data generated by HipSTR.

## RESULTS

STR genotypes defined for each individual from the newest dataset released by the 1000 Genomes Project are available in Supplementary Table 1 as an open-access database. We excluded the D21S11 marker because we did not succeed in genotyping it. Apart from this marker, the mean coverage for calling genotypes ranged from 37.14 (TPOX) to 52.53 (D12S391) (Table 1). Average successful calling rate was 98.59%, ranging from 84.18% (Penta E) to 100% (CSF1PO, D2S441, D2S1338, D3S1358, D5S818, D8S1179, D22S1045, and TPOX) (Table 2).

**Table 1.** Average coverages obtained for each STR using the HipSTR tool.

| Marker | Lowest value | Median | Highest value | Mean | Standard deviation |
| --- | --- | --- | --- | --- | --- |
| CSF1PO | 21 | 44 | 91 | 44.54 | 8.29 |
| D1S1656 | 24 | 45 | 92 | 45.48 | 8.58 |
| D2S441 | 26 | 49 | 131 | 49.17 | 9.04 |
| D2S1338 | 28 | 50 | 105 | 51.28 | 9.64 |
| D3S1358 | 28 | 51 | 119 | 51.73 | 9.33 |
| D5S818 | 20 | 42 | 98 | 42.90 | 8.14 |
| D7S820 | 20 | 38 | 86 | 39.00 | 7.78 |
| D8S1179 | 26 | 47 | 96 | 48.09 | 8.95 |
| D10S1248 | 18 | 40 | 100 | 40.75 | 7.99 |
| D12S391 | 26 | 52 | 113 | 52.53 | 9.47 |
| D13S317 | 11 | 37 | 79 | 37.93 | 7.50 |
| D16S539 | 21 | 44 | 92 | 44.70 | 8.49 |
| D18S51 | 24 | 47 | 91 | 47.38 | 9.07 |
| D19S433 | 19 | 45 | 89 | 45.28 | 8.62 |
| D22S1045 | 22 | 49 | 111 | 49.47 | 9.11 |
| FGA | 23 | 50 | 118 | 51.33 | 9.49 |
| Penta D | 19 | 43 | 95 | 43.71 | 8.74 |
| Penta E | 18 | 41 | 107 | 41.49 | 8.04 |
| TH01 | 16 | 40 | 83 | 40.73 | 8.01 |
| TPOX | 15 | 37 | 86 | 37.14 | 7.66 |
| vWA | 21 | 47 | 105 | 48.44 | 9.36 |

**Table 2.**
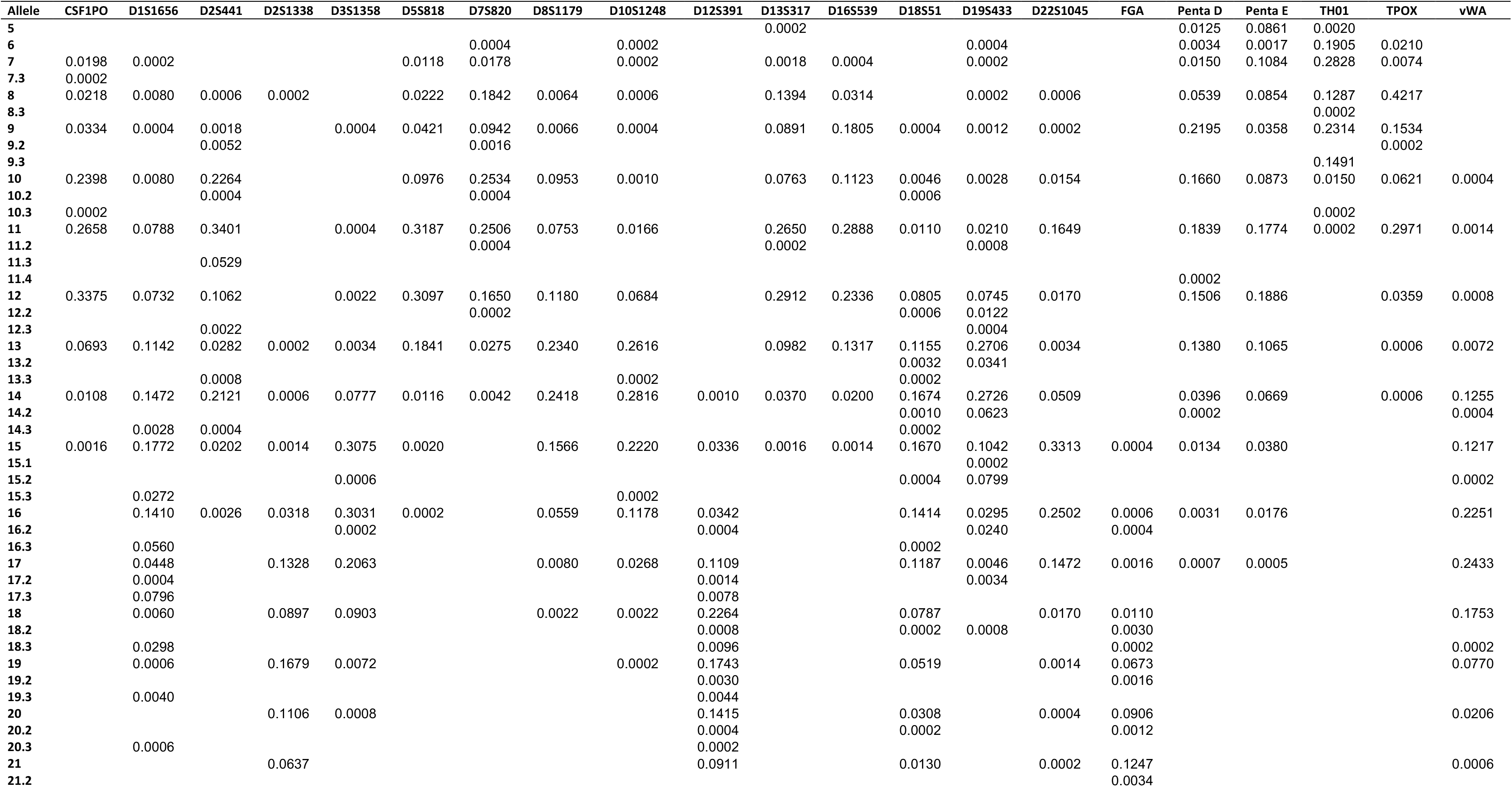
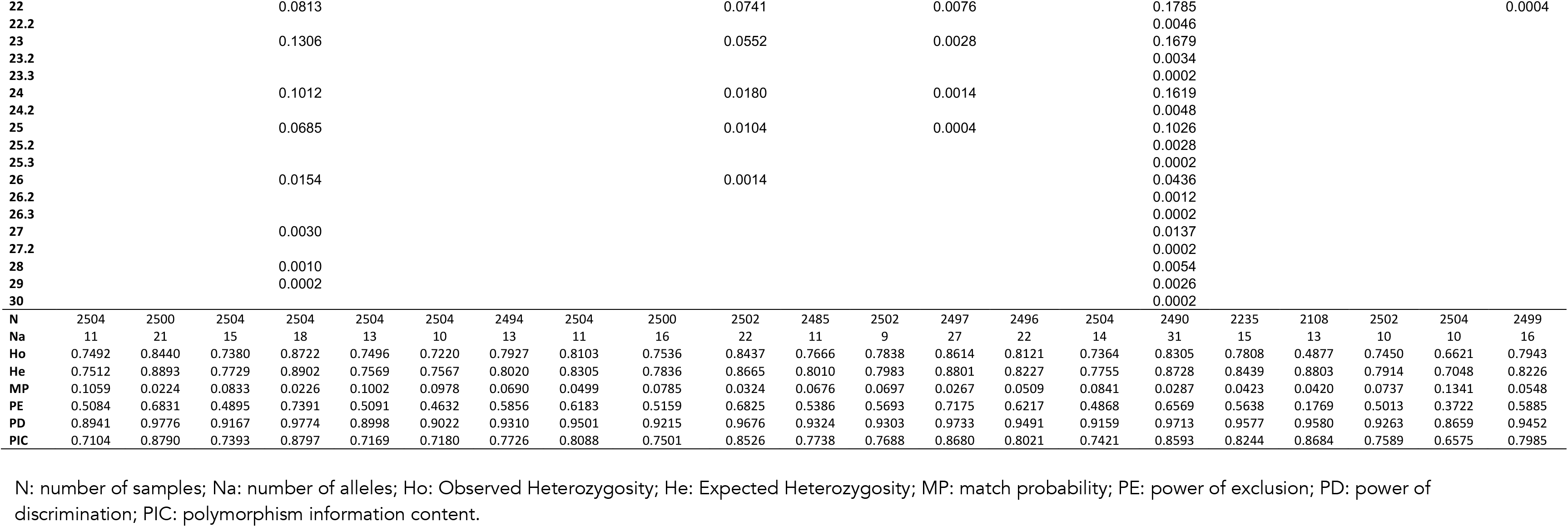
Allelic frequencies and the forensic parameters estimated for each marker in the whole 1000 Genomes dataset.

Allele frequencies and forensic parameters estimated for the whole dataset are exposed in Table 2. Allele frequencies and forensic parameters estimated for each of the 26 populations (Supplementary Table 2) and the five population groups (Supplementary Table 3) are available as supplementary data. In general, the most polymorphic *loci* in all populations were D1S1656, D2S1338, D12S391, D18S51, and FGA (Table 2). The analyzed loci were highly informative, with elevated PD values ranging between 86.59% (TPOX) and 97.76% (D1S1656). Combined MP was 5.72×10^−27^, and combined PE was 0.99999997. Analyzing each locus in each population (Supplementary Table 2), D22S1045 in PEL (71.61%) and D1S1656 in GBR (97.52%) presented the lowest and highest PD, respectively. Combined MP ranged from 1.98 × 10^−25^ in ACB to 2.20 × 10^−21^ in PEL.

Adherences of genotype frequencies to Hardy-Weinberg Equilibrium expectations were estimated for each STR at a population level (Table 3). Penta E presented heterozygote deficiency in 24 out of the 26 populations, leading to departures from the Hardy-Weinberg equilibrium. This finding clearly indicates that HipSTR incorrectly called many heterozygous genotypes as homozygous. Disregarding Penta E, the number of deviations ranged from one (D13S317 and D16S539) to five (D19S433 and Penta D), and the number of deviations across populations ranged from zero (ASW and CEU) to seven (PUR), with an average of 2.42 departures in each population. It should be emphasized that if the Bonferroni correction for multiple tests is taken into account, only 39 departures remain significant, most of them (61.53%) concerning Penta E.

**Table 3.**
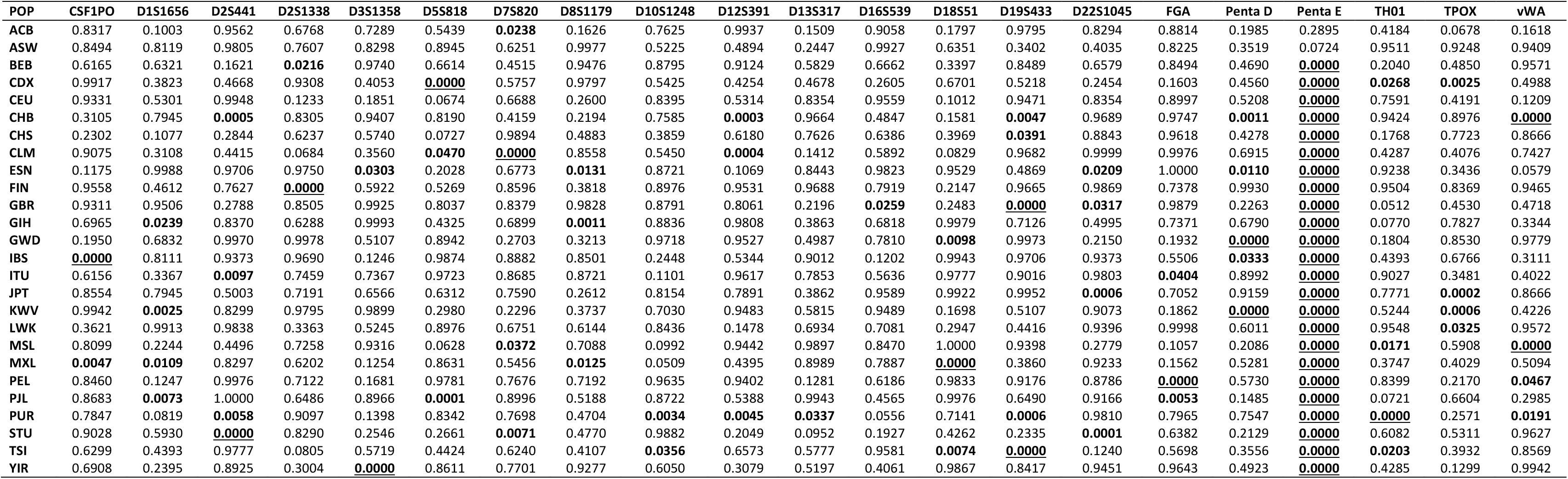
Probabilities of adherence to Hardy-Weinberg equilibrium proportions for each STR in all 26 subpopulations analyzed in the 1000 Genomes Project. Significant *p*-values (α = 0.05) are in boldface. The probabilities that remain significant after the Bonferroni correction for multiple tests (α_BONFERRONI_ = 0.05/546 = 0.000092) are also underlined.

The Principal Coordinates Analysis (PCoA) reveals four different population clusters (Figure 1). The first coordinate separated the cluster of African (AFR) populations on the right side. On the left side, it is possible to observe three different groups: the European (EUR) populations in the upper part, the East Asian (EAS) populations in the lower section, and the South Asian (SAS) populations between them. The CLM, MXL, PEL, and PUR admixed populations cluster with the European populations, while ACB and ASW cluster with Africans, reflecting their ancestry compositions.

**Figure 1.**
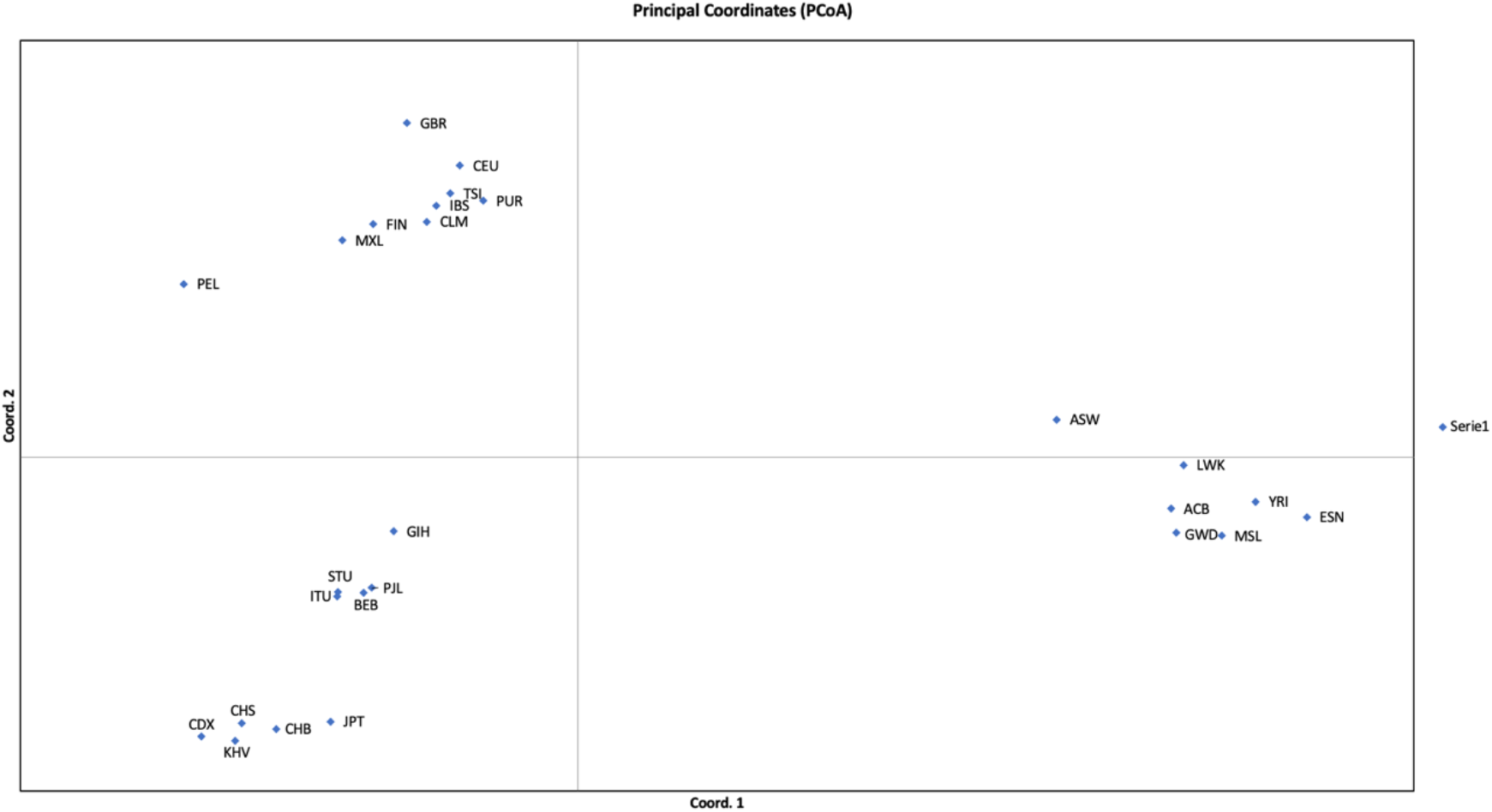
Principal Coordinates Analysis (PCoA) based on autosomal STR data from the 26 populations from the 1000 Genomes Project. Each point represents a population sample. More details on these populations are available in Supplementary Table 2. Coordinates 1 and 2 account for 39.15% and 19.25 % of the variance, respectively.

Similar results were obtained with the STRUCTURE analysis. Figure 2 depicts STRUCTURE results from runs obtained with *k* ranging from 3 to 6. With *k* = 4, each cluster reflects one of the major ancestries from the 1000 Genomes Project. Moreover, each of the six admixed American populations presents varying levels of ancestries from the four biogeographical groups.

**Figure 2.**
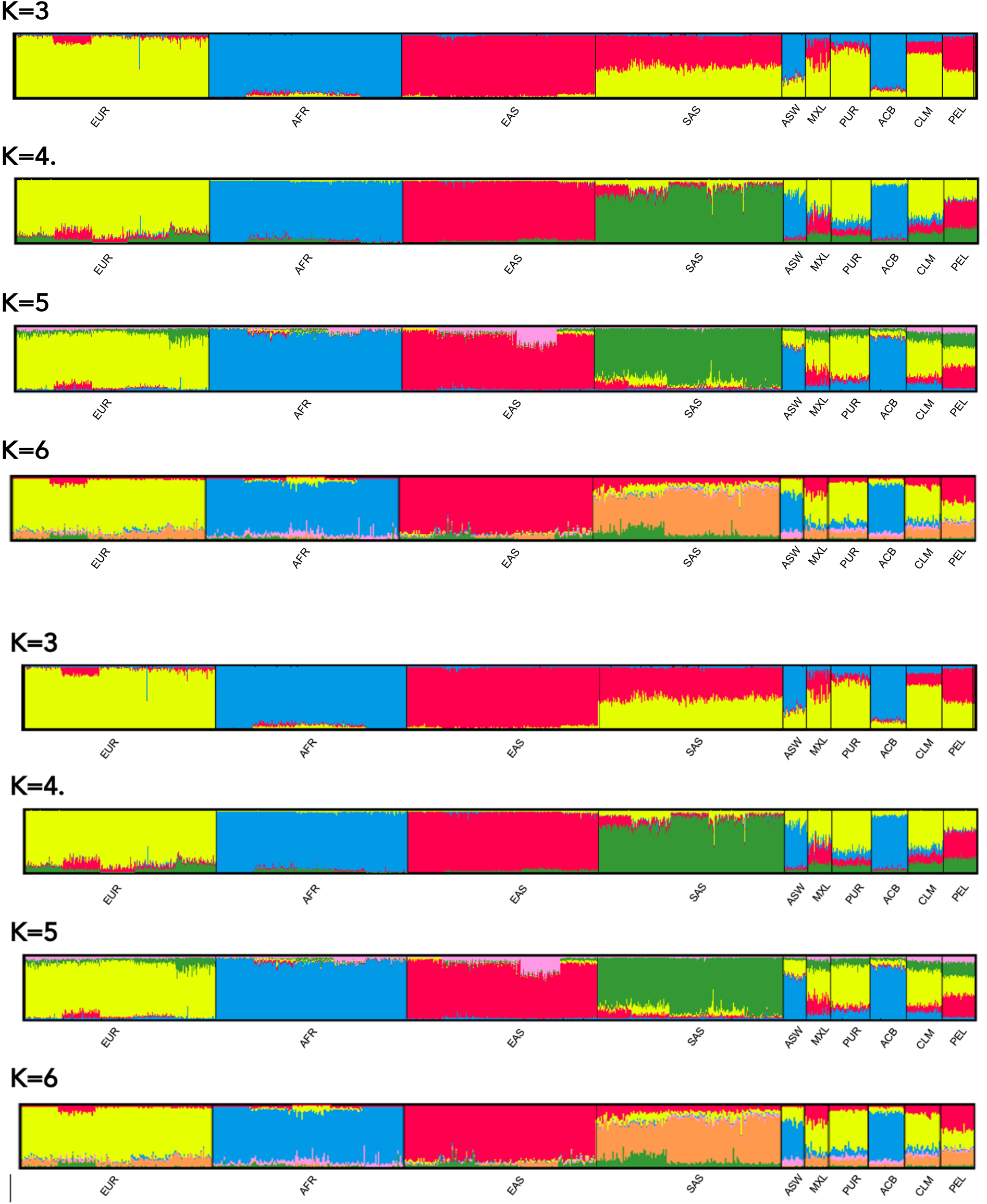
STRUCTURE analysis based on autosomal STR data from the 26 populations from the 1000 Genomes Project. Five sets of 100 independent runs, with the number of clusters ranging from 3 to 6, were conducted. Each bar plot depicts the results from the run with the largest LnP(D) for the given *k*.

To verify the distribution of variance in different levels, an AMOVA was performed assuming a hierarchical structure gathering the populations in four groups: AFR, EAS, EUR, and SAS. The six AMR populations were not taken into account. Most of the variance is observed within populations (97.12%). Differences between the four groups account for 2.54% of the variance, whereas only 0.34% of the variance occurs due to differences between populations from the same group.

Allele frequencies estimated from the 1000 Genomes dataset were also compared with STR data retrieved from for the same five major population groups (African, European, East Asian, South Asian, and admixed American) that composed the SPSmart STR browser (Pop.STR) (20) using *F*_*ST*_ (Table 4). While AMR (four), EAS (three), EUR (eight), and SAS (four) population groups presented small numbers of markers with significantly different frequencies between the two datasets, AFR presented 17 significant differences. This pattern may be reflecting the set of populations that compose the compared groups. Penta E was the only marker that showed significantly different *F*_*ST*_ values in all comparisons. Leaving AFR and Penta E aside, only 15 significant differences out of 80 comparisons are observed: the mean number of statistically significant differences was 0.75 per marker, ranging from zero (eight STRs) to three (D2S441). It should be emphasized that if the Bonferroni correction for multiple tests is taken into account, only 3 of these 15 *F*_*ST*_ values remain significant, while 6 out of 16 significant differences observed for AFR (leaving Penta E aside) and all 5 Penta E differences remain significantly different.

**Table 4.** Probabilities obtained by *F_ST_* of population differentiation comparing population groups from the 1000 Genomes Project with those from the SPSmart STR browser (Pop.STR) for each STR. Significant *p*-values (α = 0.05) are in boldface. The probabilities that remain significant after the Bonferroni correction for multiple tests (α_BONFERRONI_ = 0.05/105 = 0.00048) are also underlined.

| Marker | AFR |  | AMR |  | EAS |  | EUR |  | SAS |  |
| --- | --- | --- | --- | --- | --- | --- | --- | --- | --- | --- |
| | $F_{ST}$ | $p$-value | $F_{ST}$ | $p$-value | $F_{ST}$ | $p$-value | $F_{ST}$ | $p$-value | $F_{ST}$ | $p$-value |
| CSF1PO | 0.0084 | **0.0057±0.0007** | -0.00104 | 0.7571±0.0047 | -0.0014 | 0.6656±0.0053 | 0.0004 | 0.2445±0.0044 | 0.0019 | 0.1951±0.0036 |
| D1S1656 | 0.0024 | **0.0355±0.0019** | 0.00028 | 0.3149±0.0046 | -0.0024 | 0.9774±0.0015 | 0.0045 | **0.0000±0.0000** | 0.0021 | 0.1152±0.0032 |
| D2S441 | 0.0093 | **0.0005±0.0002** | 0.00370 | **0.0216±0.0015** | 0.0025 | 0.1251±0.0027 | 0.0074 | **0.0000±0.0000** | 0.0122 | **0.0058±0.0007** |
| D2S1338 | 0.0124 | <u>0.0000±0.0000</u> | -0.00048 | 0.6527±0.0049 | -0.0026 | 0.9940±0.0007 | 0.0031 | **0.0008±0.0003** | 0.0024 | 0.1044±0.0030 |
| D3S1358 | 0.0044 | **0.021±0.0014** | -0.00122 | 0.8369±0.0040 | 0.0010 | 0.2578±0.0044 | 0.0013 | 0.0835±0.0026 | 0.0033 | 0.1143±0.0031 |
| D5S818 | 0.0030 | **0.0477±0.0024** | 0.00609 | **0.0051±0.0008** | -0.0014 | 0.6936±0.0048 | -0.0007 | 0.7638±0.0047 | -0.0014 | 0.6250±0.0050 |
| D7S820 | 0.0047 | **0.0095±0.0009** | -0.00105 | 0.7982±0.0041 | -0.0006 | 0.5151±0.0050 | -0.0003 | 0.5677±0.0054 | -0.0023 | 0.8731±0.0034 |
| D8S1179 | 0.0033 | **0.0308±0.0018** | -0.00158 | 0.9817±0.0014 | -0.0016 | 0.7986±0.0042 | 0.0024 | **0.0150±0.0012** | -0.0003 | 0.4615±0.0049 |
| D10S1248 | 0.0017 | 0.1081±0.0033 | -0.00132 | 0.8984±0.0029 | -0.0031 | 0.9976±0.0005 | 0.0003 | 0.2627±0.0044 | -0.0001 | 0.4055±0.0046 |
| D12S391 | 0.0007 | 0.1920±0.0041 | 0.00077 | 0.1805±0.0036 | -0.0019 | 0.8774±0.0030 | 0.0008 | 0.0946±0.0030 | -0.0009 | 0.6155±0.0047 |
| D13S317 | 0.0160 | <u>0.0000±0.0000</u> | 0.00017 | 0.3478±0.0044 | -0.0018 | 0.7993±0.0045 | 0.0003 | 0.2904±0.0047 | 0.0051 | **0.0354±0.0019** |
| D16S539 | 0.0105 | <u>0.0002±0.0001</u> | -0.00051 | 0.5773±0.0047 | -0.0010 | 0.5939±0.0046 | 0.0047 | **0.0009±0.0003** | -0.0010 | 0.5987±0.0052 |
| D18S51 | 0.0018 | **0.0483±0.0019** | -0.00138 | 0.9831±0.0013 | -0.0007 | 0.5795±0.0055 | 0.0005 | 0.2028±0.0042 | -0.0003 | 0.4671±0.0047 |
| D19S433 | 0.0093 | <u>0.0003±0.0002</u> | 0.00151 | 0.1085±0.0027 | 0.0010 | 0.2476±0.0041 | 0.0000 | 0.3759±0.0049 | 0.0036 | 0.0660±0.0028 |
| D22S1045 | 0.0004 | 0.2943±0.0053 | 0.00217 | 0.0766±0.0025 | 0.0094 | **0.0055±0.0007** | 0.0034 | **0.0087±0.0009** | 0.0054 | 0.0516±0.0021 |
| FGA | 0.0011 | 0.1503±0.0037 | -0.00089 | 0.8382±0.0034 | 0.0036 | **0.0397±0.0020** | 0.0007 | 0.1306±0.0035 | 0.0018 | 0.1518±0.0039 |
| Penta D | 0.0275 | <u>0.0000±0.0000</u> | 0.00318 | **0.0113±0.0010** | -0.0015 | 0.7660±0.0042 | 0.0003 | 0.2969±0.0042 | 0.0000 | 0.4117±0.0048 |
| Penta E | 0.0139 | <u>0.0000±0.0000</u> | 0.00733 | <u>0.0000±0.0000</u> | 0.0412 | <u>0.0000±0.0000</u> | 0.0083 | <u>0.0000±0.0000</u> | 0.0179 | <u>0.0000±0.0000</u> |
| TH01 | 0.0103 | <u>0.0004±0.0002</u> | 0.00090 | 0.2083±0.0036 | -0.0025 | 0.9262±0.0027 | 0.0057 | <u>0.0001±0.0001</u> | 0.0026 | 0.1425±0.0038 |
| TPOX | 0.0039 | **0.0269±0.0019** | -0.00057 | 0.5622±0.005 | -0.0011 | 0.5341±0.0046 | 0.0015 | 0.0939±0.0033 | 0.0063 | **0.0455±0.0020** |
| vWA | 0.0065 | **0.0007±0.0003** | -0.00095 | 0.7575±0.0047 | 0.0005 | 0.3135±0.0047 | -0.0001 | 0.4312±0.0051 | -0.0010 | 0.5977±0.0049 |

## DISCUSSION

The present study provides the most diverse database of forensic autosomal STR markers from global populations. STRs display high levels of polymorphism, making them attractive for forensic purposes and population genetics studies.

Despite their usefulness in forensic genetics, STRs genotyping from NGS data, especially whole-genome sequencing assays, may be challenging. The main issues concern the high coverage required to accurate genotyping, difficulties to detect longer alleles due to reads of limited sizes, and mutations in flanking regions leading to null alleles (21). These and other problems were previously addressed by Gaag et al. (22) and Valle-Silva et al. (Valle-Silva, Frontanilla, et al., 2021, in preparation).

However, NGS technology allows the simultaneous analysis of dozens of STRs together with different classes of markers that provide complementary contributions for population genetics and human identification. For example, it is not feasible to include SNPs used as predictors of ancestry and phenotypic characteristics into commercial kits that employ capillary electrophoresis; however, it is possible to combine them with STRs in an NGS assay (1).

Despite the challenges addressed here, several studies demonstrated that it is possible to genotype STRs using dedicated bioinformatics tools. Software like LobSTR (9), toaSTR (12), STRait Razor (11), HipSTR (13), and others have shown consistent and accurate results (14)(Valle-Silva, Frontanilla, et al., 2021, in preparation). Moreover, Bornman et al. (8) demonstrated that CODIS *loci* could be accurately called even from mixtures using an NGS approach.

Willems et al. (23) presented an analysis of human STR variation using lobSTR. They used the 1000 Genomes Project phase 1 data generated using low sequencing coverage, which is excessively error-prone. In fact, they reported difficulties in detecting both alleles in each sample, resulting in an overall deficit of heterozygotes.

As previously addressed, several reasons led us to choose HipSTR to call STR genotypes from the 1000 Genomes dataset. By allowing the customization of the flanking regions, it is possible to evaluate almost any STR in hundreds of samples at once with HipSTR. It may appear more complex at first glance, but it is the most appropriate tool to deal with whole genomes. In addition, a recent evaluation of the performance of this software revealed high efficiency and accuracy levels (Valle-Silva, Frontanilla, et al., 2021, in preparation).

Despite the flexibility provided by HipSTR, the D21S11 marker could not be detected in the present study, probably because of the limited length of the Illumina NGS reads (150 bp paired-end reads) (5, 24). Gaag et al. (22), using Illumina sequencing technology, showed that longer alleles from Penta D, Penta E, and FGA presented sequencing errors at the end of the reads, resulting in null alleles and genotyping errors. Using 300 nucleotide-long paired-end reads, we successfully genotyped this marker with HipSTR (Valle-Silva, Frontanilla, et al., 2021, in preparation), which suggests a sequencing methodology issue rather than a bioinformatics one.

In this study, Penta D and Penta E showed 10.74% and 15.81% of missing data, respectively. As observed for D21S11, this problem is related to the impossibility of detecting longer alleles due to read-length constraints. Supplementary Table 4 shows the comparison between the allele frequencies estimated in the present study with those from the SPSmart STR browser (Pop.STR) (20) concerning the major groups of populations. The straightforward comparison shows that we were not able to detect alleles larger than 18 in Penta E. This failure directly led to Hardy-Weinberg equilibrium deviations (Table 3) due to deficit of heterozygotes in 24 out of the 26 studied populations. Thus, allele frequencies estimated for Penta E are strongly biased towards increased frequencies of shorter alleles and are of limited applicability (Supplementary Table 4). The probabilities obtained with the *F*_*ST*_ analysis (Table 4) support this conclusion: Penta E presented significant *F*_*ST*_ values in all five comparisons. Although Penta D and FGA also displayed this problem, their undetected alleles usually show low frequencies (Supplementary Table 4). Therefore, this technical issue did not influence the Hardy-Weinberg equilibrium and *F*_*ST*_ analysis as much as Penta E. Although this comparison is valid and helpful, it is essential to emphasize that the compared samples correspond to distinct population groups. The African population group in pop.STR comprises mainly East African Somalian individuals (404 out of 507 samples), while the 1000 Genomes Project samples corresponded to West Africa. Similarly, more than 50% of the European population group from pop.STR is composed mainly of U.S. Europeans (1443 out of 2135) (5,20). Taken together, these results support the robustness of the bioinformatics analysis performed in the present study and the reliability of allele frequencies distributions for all loci except for Penta E.

The most polymorphic *loci* in the whole 1000 Genomes dataset were D1S1656, D2S1338, D12S391, D18S51, and FGA. All these markers presented high degrees of polymorphism throughout the world. AMOVA revealed that most of the variance (97.12%) in allele frequencies is observed within populations, corroborating previous studies (25, 26). However, as expected, the principal component analysis (Figure 1), the clustering analysis performed with STRUCTURE (Figure 2), and the AMOVA confirmed that the four ancestral populations groups (AFR, EUR, EAS, SAS) defined by the 1000 Genomes Consortium are indeed significantly different from each other. Since the admixed American populations present different ancestry compositions (Figure 2), most of them cluster with Europeans, while ACB and ASW cluster with Africans (Figure 1).

The results obtained with the STRUCTURE software corroborate the relationship between the different population groups and provide additional support for the reliability of the genotypes calculated. If *k* = 3, SAS resembles an admixture between EAS and EUR. A specific cluster for SAS emerges with k = 4. With *k* = 5, a minor Eurasian (shared between EUR and EAS) component arises. With *k* = 6, the SAS shared ancestry with EUR and EAS becomes more evident. Regarding the admixed American populations, irrespective of the number of clusters considered, ACB and ASW reveal their preeminent African origin, CLM and PUR reveal more extensive European ancestry, and MXL and PEL reveal almost equal amounts of European and Amerindian (i.e., EAS) ancestries. These results fully corroborate the distribution of the populations into the PCoA (Figure 1). Additional clusters do not provide increased resolution with straightforward meaning (data not shown).

In conclusion, this investigation offers a reliable open-access STR database from the 1000 Genomes Project Phase 3 high-coverage (30x) WGS data generated by the NYGC. However, the limited length of sequencing reads introduced noticeable bias in allele frequencies estimated for Penta D and Penta E. The reliability of this dataset is supported (a) by previous studies that attested HipSTR efficiency, (b) by the Hardy-Weinberg equilibrium analysis, (c) by the set of analyses employed to evaluate interpopulation genetic diversity population, and (d) by the comparison of the allele frequencies obtained here with those of other initiatives that used capillary electrophoresis. Despite our eagerness that this open-access database will be of great interest for future forensic population genetics studies, it is imperative to highlight that the current 1000 Genomes Project dataset does not describe the worldwide human genetic diversity. In fact, many biogeographical regions, mainly from Oceania and the Americas, were not sampled, indicating that additional large-scale initiatives may provide further insight into worldwide STR population diversity.

## ACKNOWLEDGEMENTS

We thank Dr. Thomas Willems for sharing his knowledge and expertise in bioinformatics and population genetics and his support with the HipSTR tool. This study was financed in part by the Coordenação de Aperfeiçoamento de Pessoal de Nível Superior - Brasil (CAPES) - Finance Code 001. C.T.M.J. (#312802/2018-8) is supported by a Research fellowship from CNPq/Brazil.

## Notes

https://drive.google.com/drive/folders/1CKNMBNTKXJ08qzUqj9YKt2RauYfTyHUk?usp=sharing

### Competing Interest Statement

The authors have declared no competing interest.

